# Extracting deep learning based morphology segmentation footprint for boar sperm cells

**DOI:** 10.64898/2026.08.07.743571

**Authors:** Joonhyung Park, Megan Ratka, Angona Biswas, Ian Shofner, Karl Kerns, Anwesha Sarkar

## Abstract

Reliable delineation of the head and tail of swine spermatozoa supports automated assessment of boar semen quality, from morphometric measurement to the quality control of insemination doses. In practice this relies on fluorescent staining, which adds chemistry, cost, and delay to every acquisition and labels only the nucleus. Recent work coupling imaging flow cytometry with machine learning has advanced rapidly, yet the segmentation stage still depends on a stained channel at inference and resolves the head alone. We present a supervised encoder decoder network that segments boar spermatozoa from brightfield images acquired on an Amnis ImageStream Mark II with no stain at inference. Training labels derive from the Hoechst 33342 nuclear channel (Ch7), recorded in registration with brightfield (Ch1); the dye serves only as an annotation source, and the network sees Ch1 alone. The best semantic segmentation model reaches a Dice coefficient of 0.940 on held-out cells. For comparison we evaluate a classical morphological pipeline, four further semantic segmentation models spanning three decoder families and two ImageNet-pretrained backbones, and two zero-shot pipelines built on the Segment Anything Model 2 (SAM 2), prompted either by a dilated box around the predicted head mask or by head and tail boxes emitted by a Gemma 4 Vision Language Model (VLM). The zero-shot route scores 0.637 against Ch7 but labels the tail, which the fluorescence protocol cannot. Cells scoring worst under the supervised model proved to be mostly registration failures rather than segmentation failures, as Ch7 is displaced relative to Ch1. Manual screening for this drift is infeasible at dataset scale, so we propose a flagging system that marks any Dice below 0.792, two standard deviations below the mean, and pairs it with a zero-shot pipeline in which a VLM l and SAM 2 cross-check the flagged cell before human review.

## 1 Introduction

Morphological assessment of boar spermatozoa is a routine step in the quality control of semen intended for insemination, and it rests on delineating the head and the tail of individual cells. Head dimensions, tail defects, and the classification of a cell as normal or aberrant all follow from that delineation. In current practice it is obtained through fluorescent staining, which resolves the nucleus but leaves the flagellum unlabeled and adds hours of sample preparation to a workflow constrained by the viability of the sample. Manual tracing avoids the dye but is subjective and does not scale to the hundreds of thousands of cells in a single insemination dose. Automated, stain-free segmentation is therefore the enabling step for morphological phenotyping at scale, and it is the problem this work addresses. The mammalian spermatozoon is a polarized cell whose principal regions are the head, the midpiece, and the tail, each carrying distinct diagnostic weight in fertility assessment [1]

(Fig. 1). The head contains the condensed nucleus and the overlying acrosome. In the boar it is approximately elliptical in projection, with reported population means near 8.29 µm in length, 4.26 µm in width, and 27.79 µm^2^ in projected area [2]; this work returns to that elliptical assumption in Section 2.3.2. Acrosomal integrity is a routine predictor of boar fertility [3], but its boundaries resist clean resolution under conventional stains [4] and are not recovered here.

**Figure 1:**
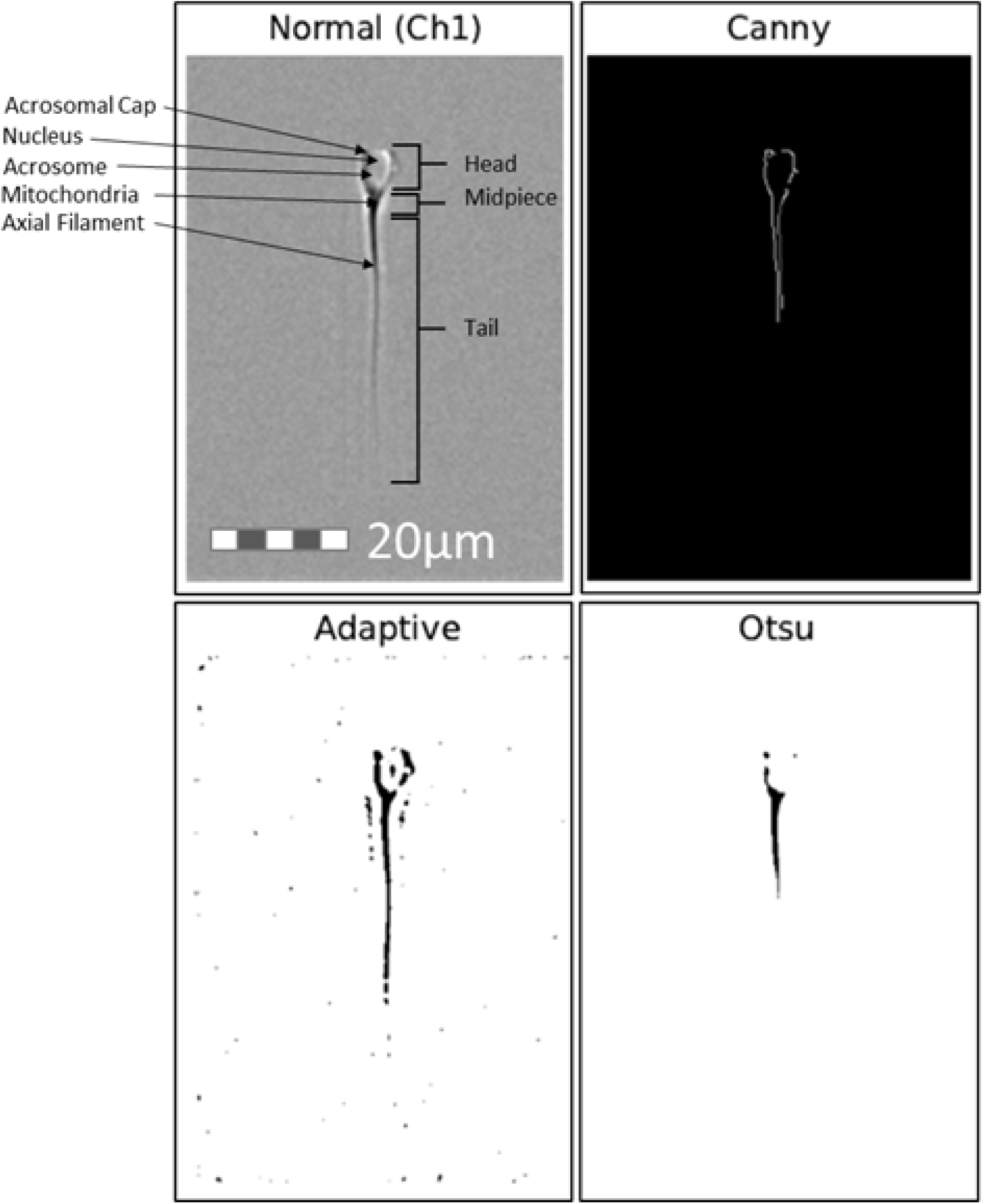
Sperm anatomy and classical computer-vision operators on a Ch1 brightfield cell. Top left: the raw grayscale image, annotated with the acrosomal cap, nucleus, acrosome, mitochondria, and axial filament, and with the head, midpiece, and tail marked. Scale bar 20 µm. Top right: Canny edge detection. Bottom left: adaptive thresholding. Bottom right: Otsu’s global thresholding.

The midpiece connects the head to the flagellum and houses the mitochondrial sheath that powers motility. Its boundaries, like those of the acrosome, resist delimitation by dye alone. The tail is the motile apparatus of the cell, and its dimensions, together with those of the head and midpiece, jointly determine swimming behavior [5]. This work segments the head and the tail; the midpiece is not recovered, although its morphometry is of direct interest, and Section 4.6 returns to it. Real boar samples contain tens of thousands of cells that depart from this idealized description, and any automated method must accommodate that variation. The departures are not arbitrary: a small number recur often enough, and carry enough diagnostic weight, that clinical assessment treats them as distinct morphological classes [4]. Their consequences for segmentation are geometric. A cytoplasmic droplet distorts the outline at the neck, a strongly coiled tail overlaps the head in projection, a ruffled acrosome opens the head boundary that a smooth oval model assumes, and a cell rotated out of the imaging plane presents no measurable outline at all. Each of these alters the shape, the local contrast, or both, so no single set of fixed parameters covers them. The classical operators evaluated in Section 2.3 are brittle on this data for that reason, and per-pixel learned prediction is more robust to it. The networks here are trained on every Ch1/Ch7 pair that passes quality control, without stratification by morphology, so the reported scores are population means over a heterogeneous sample; Section 4.6 returns to this.

The data in this work were acquired with a Cytek Amnis ImageStreamX Mark II imaging flow cytometer [6], an instrument that combines the throughput of flow cytometry with the spatial resolution of a microscope. As cells traverse the fluidic stream, the instrument captures a multi-channel image stack of each event, writing one image per channel. The instrument provides 12 channels across two cameras, Ch1 through Ch6 on Camera 1 and Ch7 through Ch12 on Camera 2. Two are used here: channel 1 (Ch1), a brightfield image of the whole cell, and channel 7 (Ch7), a fluorescence image in which Hoechst 33342 binds the condensed sperm nucleus and so marks the head [4]. Ch1 is the sole model input throughout this work. Ch7 supplies the head annotation, and Section 2.2 defines how it is thresholded into a per-pixel label.

Recovering the head from Ch1 alone is not a new problem, and the classical approach to it is well established. Automated cell-morphology analysis has long relied on hand-designed operators: edge detectors such as Canny, global and adaptive thresholding schemes including the Otsu method [7], and morphological filtering pairs like opening and closing [8]. Fig. 1 shows three of these applied to a Ch1 brightfield cell. Each recovers part of the outline and none recovers all of it: edges break where contrast falls off along the tail, and a single global threshold cannot hold for both the dense head and the thin flagellum.

Such operators are computationally cheap and their behavior is interpretable. Preliminary experiments in the present study established that Otsu thresholding combined with active contours, morphological closing, or custom erosion–dilation sweeps could recover the coarse boundary of an uncrowded cell, in line with earlier work segmenting the sperm head, acrosome, and nucleus in human semen smears [9]. The difficulty is that these operators are fragile at scale. They are sensitive to sensor noise and focal-plane drift, and above all to the morphological variation described above. A parameter configuration that isolates the head of one cell fails on the next, and no single choice of kernel size, structuring element, or threshold value covers the full morphological range of the sample. Section 3.1 quantifies this ceiling.

Modern semantic-segmentation architectures address these constraints by replacing rigid, handcrafted kernels with self-learned, per-pixel predictors. Encoder–decoder structures such as U-Net [10], together with nested variants like UNet++ [11] and multi-scale attention networks like MA-Net [12], were developed for precisely this class of problem: recovering boundaries that vary in shape and contrast from one instance to the next. Complementing these supervised frameworks, foundation models such as the Segment Anything Model (SAM) [13] introduce zero-shot, promptable segmentation driven by bounding-box or centroid prompts, and when coupled with vision–language models they can be directed toward a named structure without task-specific training. The practical advantage over classical operators is stability: where a threshold or structuring element must be re-tuned from one cell to the next, learned models derive their decision rules from data and hold across morphological variation. What remains open for the present application is whether that stability survives the transfer from a stained channel to an unstained one, and whether it extends to the flagellum, which no fluorescence protocol labels.

A central practical motivation for this work is the elimination of chemical dyes from routine morphological analysis. In a production andrology laboratory, freshly collected boar semen must ideally be received, analyzed, and processed into individual insemination doses within an hour or less, yet fluorescence-based biomarker assays are poorly suited to that timeline: a single stained assay can require between roughly 1.5 and 8 hours of preparation and incubation before analysis, and a complete diagnostic workup typically stacks several such assays in series. Beyond time, fluorescence workflows carry a substantial resource burden. Fluorescent-based analysis equipment and their associated optics are capital-intensive instruments, the per-sample cost of probes and consumables is high, and each stained assay demands additional skilled labor for preparation, compensation, gating, and analysis. A method that recovers head and tail morphology directly from label-free brightfield images therefore removes a genuine throughput and cost bottleneck, allowing morphological phenotyping to keep pace with the sub-hour processing window that commercial semen handling demands while reserving expensive fluorescence chemistry for the confirmatory assays that truly require it. It also lowers the barrier to entry for the wider research community, for whom the equipment and per-sample expense of fluorescence imaging can be prohibitive, and it establishes the foundation for stain-free morphological phenotyping across other mammalian species.

In this paper we present a comparative study of classical, supervised, and foundation-model approaches to head and tail segmentation of boar spermatozoa, and combine the strongest of them into a stain-free pipeline in which the best semantic segmentation model delineates the head and a Gemma-4 26B MoE vision–language model coupled with SAM 2 delineates both the head and the tail. Because the two routes are independent, their disagreement on a given cell identifies off-centered Ch1/Ch7 pairs and flags misalignment between the two channels. Our key contributions are as follows:

- **Paired imaging dataset and misalignment protocol:** We assemble 11,732 co-registered Ch1/Ch7 boar sperm image pairs across five acquisition folders. A separate 2,172-pair set, annotated for channel alignment, trains a model used solely to screen the full dataset; it identifies 312 misaligned pairs, leaving 11,420 clean pairs and a reproducible protocol for detecting inter-channel drift at scale.
- **Classical segmentation baseline:** We evaluate a family of handcrafted, non-learned pipelines, including morphological erode–dilate–erode parameter sweeps, a four-stage local-contrast head-hunting pipeline, and an elliptical-stability analysis of the head aspect ratio. These characterize the accuracy ceiling of fixed-parameter operators on morphologically heterogeneous data.
- **Stain-free supervised segmentation:** We compare five encoder–decoder models and establish UNet++ with a ResNet-50 encoder as the strongest, reaching a Dice coefficient of 0.940 from brightfield input alone. We show that preserving native input resolution by zero-padding (165 × 120 to 192 × 128) rather than downscaling to a power-of-two grid yields (165 × 120 to 128 × 96) a +2.5-point absolute gain, while 4× ESRGAN super-resolution preprocessing contributes approximately 0.0025, indicating the limits of generative upscaling on subcellular microscopy data.
- **Evaluation of zero-shot foundation models:** We assess bounding-box-prompted SAM 1 (ViT-H) and SAM 2 (hiera-large) as head refinement stages, using a prompt taken from the mask predicted by the supervised model. The average head score against Ch7 falls below the supervised baseline, but the average is misleading: the model usually either finds the head well or misses it entirely, with little in between. The failures fit what is expected from a model trained on natural images and prompted with a coarse box on an object of roughly 150 pixels, and super-resolution does not help.
- **Tail recovery and misalignment flagging:** Prompted by a Gemma-4 26B MoE vision–language model instead of by a predicted mask, SAM 2 delineates both the head and the tail. The tail is not labeled by the fluorescence protocol, so the supervised route cannot learn it. Because this route uses neither the network nor the dye, it gives a second opinion on every cell: where the two routes agree, the mask can be trusted, and where they disagree, the Ch1/Ch7 pair is often misaligned. We use this to flag cells for human review in a dataset too large to check by eye.

The remainder of this paper is organized as follows. Section 2 describes the imaging protocol, the construction of the per-pixel head label from the Ch7 fluorescence channel, and the four experimental routes: classical operators, supervised networks, box-prompted foundation models, and the vision–language pipeline. Section 3 reports the quantitative outcome of each route together with the channel-misalignment analysis. Section 4 interprets those results, sets out the proposed routing scheme, and states the limitations of the study. Section 5 concludes.

## 2 Methods

This section describes the experimental setup and the mathematical formulations used throughout. The experimental stages are presented in the order they were carried out, since the result of each shaped the design of the next. We begin with the Ch7 fluorescence channel and the per-pixel head labels derived from it, which serve as the reference throughout, and close with the evaluation protocol common to all stages.

### 2.1 Sample Preparation and Imaging Flow Cytometry

The image data analyzed in this study was generated following the boar semen preparation and imaging flow cytometry protocols established for our prior porcine sperm studies [14, 15]. Semen was collected from commercial Duroc boars of known fertility, housed one boar per pen at a single commercial boar stud and maintained on a standard commercial boar diet, using the two-gloved-hand technique. Only ejaculates exhibiting at least 80% progressive motility were retained. Immediately after collection, each ejaculate was extended five-fold in Preserve Xtreme extender held within 2 °C of the semen and transported to Iowa State University the same day, where it was recorded and stored at 17 °C. Semen used in this study was excess from industry production and not directly collected for this study, thus exempt from Institutional Animal Care & Use Committee (IACUC) oversight. Within 24 h of collection, sperm concentration and motility were confirmed by computer-assisted semen analysis (CASA) and each sample was diluted to a working concentration of 10 million sperm per mL.

For CASA, a 1000 µL aliquot was equilibrated at 37 °C for 10 min, mixed gently to distribute cells evenly, and 3–3.5 µL was loaded into a 20 µm disposable counting chamber (Minitübe GmbH, Tiefenbach, Germany). Concentration and motility were quantified on a Zeiss Axioscope 5 microscope fitted with a Basler ace acA2440-75uc camera and a 10×/0.25 A-Plan objective using Minitube AndroVision software. For imaging, a 5-million-cell aliquot was centrifuged (110 × g, 5 min) and resuspended in 100 µL of porcine non-capacitation medium containing Hoechst 33342 (H33342; 1:1000), which labels the condensed sperm nucleus and thereby designates the head region. Samples were incubated for 30 min at room temperature in the dark, washed by centrifugation, and resuspended in phosphate-buffered saline.

Image acquisition was performed on a Cytek Amnis ImageStream Mark II imaging flow cytometer (Fremont, CA, USA) fitted with a 40× objective and operated at a flow-core diameter and speed of 6 µm and 66 mm/s, with SpeedBeads used to maintain focus. Raw multi-channel image stacks were collected in INSPIRE software (v3.0), recording a brightfield image on channel 1 (Ch1) and the Hoechst 33342 nuclear image on channel 7 (Ch7), with the 405 nm laser (10 mW) exciting H33342. Events were compensated and gated for focus and single cells in IDEAS software (v6.4), and the Feature Finder function was used to exclude spermatozoa laterally aligned to the camera so that retained events present the head and tail within the imaging plane. The spatially registered Ch1/Ch7 image pairs produced by this workflow constitute the paired brightfield/fluorescence dataset used throughout this study: for the present segmentation task only these two channels are retained, Ch1 supplies the sole model input, and the thresholded Ch7 nucleus supplies the head ground truth, as defined in the following subsection. Images were exported from IDEAS software to TIFF format for the experiments.

### 2.2 Ground-Truth Definition

#### 2.2.1 Head Label via Fluorescence Thresholding

Because Ch7 fluorescence is spatially confined to the head, a thresholded Ch7 image constitutes a per-pixel head label. A pixel was assigned to the head class where its Ch7 intensity satisfied

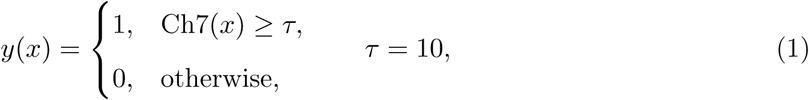

where x indexes pixel location and y(x) is the binary head label. The threshold was set by inspecting the Ch7 intensity distribution. The signal is extremely sparse: approximately 99% of pixels are exactly zero, with a thin residual population of weak values in the 1–9 range attributable to background autofluorescence. A threshold of τ = 10 excludes this residual population while preserving faint head-boundary pixels. The choice biases the label in a controlled direction, since lower thresholds enlarge the head region and favor recall while higher thresholds contract the boundary and favor precision.

Over the range [7, 10] the mask changes by a mean of 2 pixels, roughly 1% of its area (Fig. 2), consistent with the Ch7 image being very nearly binary in its spatial support. The same fixed threshold was applied to every image in every acquisition folder, so the label rule is identical across the dataset.

**Figure 2:**
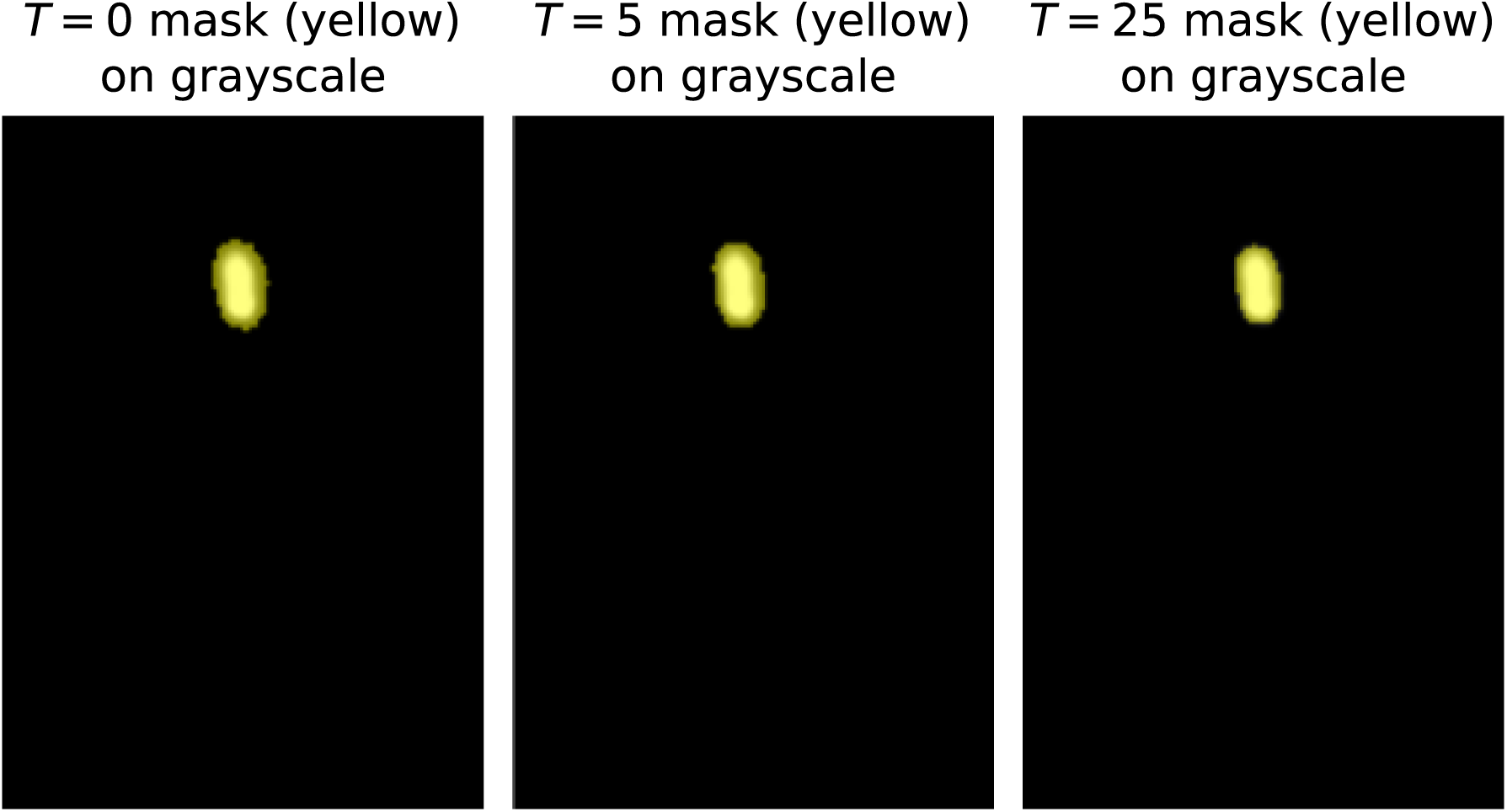
Effect of the intensity threshold τ on Ch7 head-mask extent. Lower τ favors boundary recall; higher τ favors precision.

#### 2.2.2 The Fluorescence-Bloom Observation

To compare the two channels, the binarized Ch7 mask was overlaid on the corresponding Ch1 brightfield image at 50% transparency. The overlay showed a clear discrepancy: the fluorescence-defined head region is consistently larger than the head visible in brightfield. Two explanations are available, and both may contribute. The first is the brightfield image itself, whose intensities occupy a narrow 8-bit band peaked in the 156–166 range, making the true cell border hard to localize. The second is the fluorophore, whose emission spreads optically beyond the physical nucleus, so that any nonzero-intensity cutoff tends to overstate the head’s extent. Either way, the fluorescence boundary and the visible brightfield boundary do not coincide.

The Ch7 mask is therefore treated as an operational reference rather than as anatomical truth. It is reproducible, it follows a stated rule, and it is the only head label available at this scale, but it locates the boundary where the fluorophore emits rather than where the cell edge lies. Every Dice figure reported in this work is scored against that reference, and Section 4.6 returns to what it does and does not establish. The distinction also anticipates a result in Section 2.5: models trained on natural images segment toward what is visually salient rather than toward where the fluorophore binds, so they are penalized by a reference that does not mark the same boundary they find.

#### 2.2.3 Absence of Tail Ground Truth

No channel in the present acquisition protocol labels the tail. The tail is resolved in Ch1 but is unmarked in Ch7 and in every other collected channel. This is a property of the stain rather than of the acquisition: Hoechst 33342 binds DNA, which in the mature spermatozoon is confined to the condensed nucleus, so the flagellum carries no signal to threshold. Supervised training is therefore structurally restricted to the head, and recovery of the tail requires either manual annotation or labels derived from an auxiliary source. This asymmetry motivates the foundation-model experiments of Section 2.5 and the vision–language-model route of Section 2.6.

### 2.3 Classical Segmentation Baseline

Before any learned model was trained, a family of handcrafted operators was evaluated to establish the accuracy attainable without learning and to characterize the geometry of the Ch7 reference itself.

#### 2.3.1 Erode–Dilate–Erode Grid Sweep

Fixed morphological schedules did not separate head from tail reliably across the sample, so a systematic sweep was run over a three-stage erode–dilate–erode cascade, parameterized jointly by an adaptive intensity threshold T ∈ [0, 40] and a per-stage iteration count M ∈ [1, 7]. The sweep used repeated iterations of a 3 × 3 structuring element, exploiting the associative property of Minkowski addition [16, 17]: n sequential operations with a 3 × 3 square kernel are equivalent to a single operation with a (2n+ 1) × (2n+ 1) kernel [18], which allows the grid search to reach large receptive fields at low cost. The best configuration reached a Dice score of 0.551. The highest-scoring masks were not the most faithful ones: peak Dice was often attained by bloated, over-dilated regions rather than well-fitted boundaries, indicating that the cascade dilates past the true edge rather than settling onto it.

#### 2.3.2 Elliptical-Stability Analysis of the Aspect Ratio

To characterize the head’s geometry independently of any single threshold, the binarized Ch7 region was fitted with an ellipse across the full 8-bit intensity range, T ∈ {0, 1, . . ., 255}. For each T the semi-major (a) and semi-minor (b) axes were recorded, giving the area

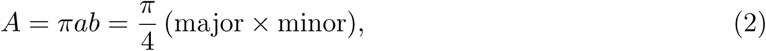

and the aspect ratio b/a ∈ (0, 1]. Tracking b/a against T tests whether threshold selection distorts the fitted shape. The aspect ratio declined from roughly 0.60 at T = 0 to 0.38 at high thresholds, so the choice of threshold is consequential and must be bounded. A 3σ envelope of the aspect ratio was therefore adopted, admitting 99.8% of the observed spread as valid for Ch1 matching. Two independent estimates converged: a full threshold-sweep average and a per-cell Otsu threshold gave aspect-ratio means of 0.471 and 0.461.

#### 2.3.3 Four-Stage Head-Hunting Pipeline

The sweep results were consolidated into a four-stage pipeline that localizes the head rather than the tail, on the premise that once the head boundary is excluded from the whole-cell contour, the tail is recovered as the residual at no additional cost.

The first stage isolates the cell from the background by combining a local-contrast operator with an adaptive intensity criterion. Local contrast is the intensity range within a 3 × 3 neighborhood, C(x, y) = max_(3*×*3)_(I) − min_(3*×*3)_(I), and a pixel is admitted to the cell mask only if it is both darker than the local mean and situated in a region of sufficient contrast,

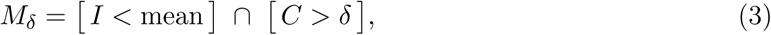

where δ is a sensitivity threshold. The first condition exploits the cell being darker than its surround under brightfield illumination; the second rejects the flat, low-contrast background.

The second stage consolidates the fragmented mask M*_δ_* into a single filled region by morphological closing, retention of the largest connected component, and interior filling. The structuring element was swept from 3 × 3 to 11 × 11 and locked at 5 × 5; on a 165 × 120 image, kernels above 11 × 11 dilate the boundary past any faithful representation of the contour.

The third stage localizes the head by fitting the largest admissible ellipse lying substantially within the cell. For a candidate ellipse E, constrained to the 3σ aspect-ratio envelope of Section 2.3.2, coverage is the fraction falling inside the cell region S, and the selected estimate is

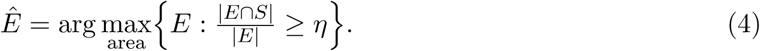

The acceptance threshold was swept over η ∈ [0.6, 0.95] and locked at 0.85, admitting the largest safe ellipse while rejecting candidates that protrude appreciably beyond the cell.

The fourth stage scores a detection by combining a distributional overlap term with a detection-rate term. The overlap term is the overlapping coefficient between the aspect-ratio distribution recovered from Ch1 and the reference distribution from Ch7, bounded in [0, 1]; multiplying by the detection rate penalizes configurations that match the reference distribution on only a small fraction of cells. The criterion was validated on a held-out set of 2,346 images against both aspect ratio and area.

### 2.4 Supervised Semantic Segmentation

#### 2.4.1 Architectures

Five encoder–decoder models were trained in PyTorch, spanning three decoder families (U-Net, UNet++, MA-Net) and two encoder backbones (ResNet-50, EfficientNet-B3), each carrying ImageNet-pretrained weights (Table 1). In every case the encoder is initialized from its pretrained state and the decoder from scratch; skip connections carry features from each encoder stage to the corresponding decoder stage, letting the decoder reconstruct a pixel-accurate mask from semantically rich but spatially coarse encoder representations.

**Table 1:** Five encoder–decoder configurations. All encoders carry weights pretrained on ImageNet.

| # | Decoder | Encoder | Pretrained |
| --- | --- | --- | --- |
| 1 | U-Net | ResNet-50 | ImageNet |
| 2 | U-Net | EfficientNet-B3 | ImageNet |
| 3 | UNet++ | ResNet-50 | ImageNet |
| 4 | UNet++ | EfficientNet-B3 | ImageNet |
| 5 | MA-Net | EfficientNet-B3 | ImageNet |

#### 2.4.2 Loss Function

Training minimized a composite objective summing a per-pixel binary cross-entropy term and a soft Dice term [8, 10],

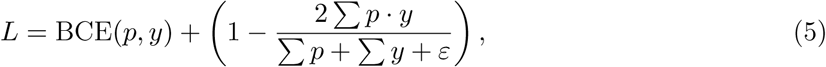

where p is the predicted per-pixel probability, y is the binary label of Equation 1, the sums run over all pixels, and ε ≈ 10*^−^*^8^ prevents division by zero. The pairing is deliberate: cross-entropy supplies smooth per-pixel gradients that calibrate the predicted probabilities, while the soft Dice term confers robustness to the foreground–background imbalance intrinsic to this task, in which head pixels constitute roughly 0.75% of the image [4, 9]. A pure cross-entropy objective would be dominated by the background majority; the Dice term restores sensitivity to the minority foreground.

#### 2.4.3 Training Configuration

Models were optimized with Adam at an initial learning rate of 10*^−^*^4^ under a ReduceLROnPlateau schedule (factor 0.5, patience 5 epochs) and a batch size of 16. An exploratory model trained on an independent set of 1,737 pairs ran for 50 epochs; the four full-dataset models ran for 80.

Augmentation was applied to encourage invariance to geometric transformation [19]. Cells appear at arbitrary orientation in the imaging plane, so random horizontal and vertical flips (p = 0.5) and uniform rotation over [−180*^◦^*, 180*^◦^*] (p = 0.8) discourage reliance on absolute orientation and promote orientation-independent features [10, 19]. To counteract variation in exposure, stain density, and sensor artifacts across imaging sessions, random brightness and contrast perturbations (p = 0.4) and additive Gaussian noise (p = 0.3) decouple structural boundaries from local illumination gradients [10], preventing the network from relying on low-level intensity statistics to locate shapes [20]. An elastic-deformation transform used in earlier iterations was removed: elastic warping bends the thin tail into anatomically implausible shapes, adding variance that impairs rather than improves generalization.

#### 2.4.4 Input Sizing: Resize versus Pad

The native resolution of 165 × 120 is not divisible by 32, the downsampling factor imposed by the ResNet-50 encoder’s five stages. Two strategies were compared under otherwise identical conditions (hyperparameters, augmentations, data split, random seed):

- **Resize:** downscale to 128 × 96, preserving aspect ratio but discarding roughly 21% of the native pixels.
- **Pad:** zero-pad the bottom and right margins to 192 × 128, preserving every native pixel at the cost of empty borders.

All five models were trained under both strategies and evaluated at the native 165 × 120 resolution, with resize predictions upsampled back and pad predictions cropped to the native region. The pad strategy outperformed resizing across all five, giving a +2.5-point absolute gain in Dice for the best model (UNet++ with ResNet-50), and the 192 × 128 configuration was adopted.

The likely reason lies in the head’s scale. The head spans roughly 12–15 pixels at native resolution, and the fluorescence bloom places the reference boundary a fixed one to three pixels outside the visible edge. Downscaling by a linear factor of 0.78 shrinks the head to roughly 9–12 pixels, so that fixed offset occupies a proportionally larger fraction of the object. Because Dice is a normalized overlap measure, a fixed absolute error inflates in relative terms as the object shrinks.

### 2.5 Zero-Shot Foundation Models

#### 2.5.1 SAM Refinement of Predicted Head Masks

With a supervised baseline established, the study tested whether the Segment Anything Model could refine the best UNet++ head predictions in a zero-shot setting. Two variants were evaluated: SAM 1 with the ViT-H backbone and SAM 2 with the hiera-large backbone. The pipeline proceeds in four steps. The UNet++/ResNet-50 model produces a head mask; a tight axis-aligned bounding box is computed around it; each side is extended outward by 30% of its width or height, with clipping to the image bounds, giving the model tolerance in case the predicted box is marginally too tight,

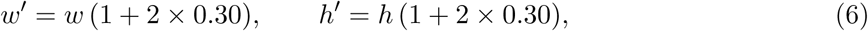

where w, h are the tight dimensions and w*^′^*, h*^′^* the padded ones; and the full Ch1 image is passed to the model with the padded box as a single-mask prompt, returning one binary mask at native resolution (Fig. 3). Both variants received identical box prompts. No point or text prompts were supplied, and neither model was fine-tuned on the present data, so the experiment measures how well natural-image foundation models transfer zero-shot to fluorescence-defined microscopy.

**Figure 3:**
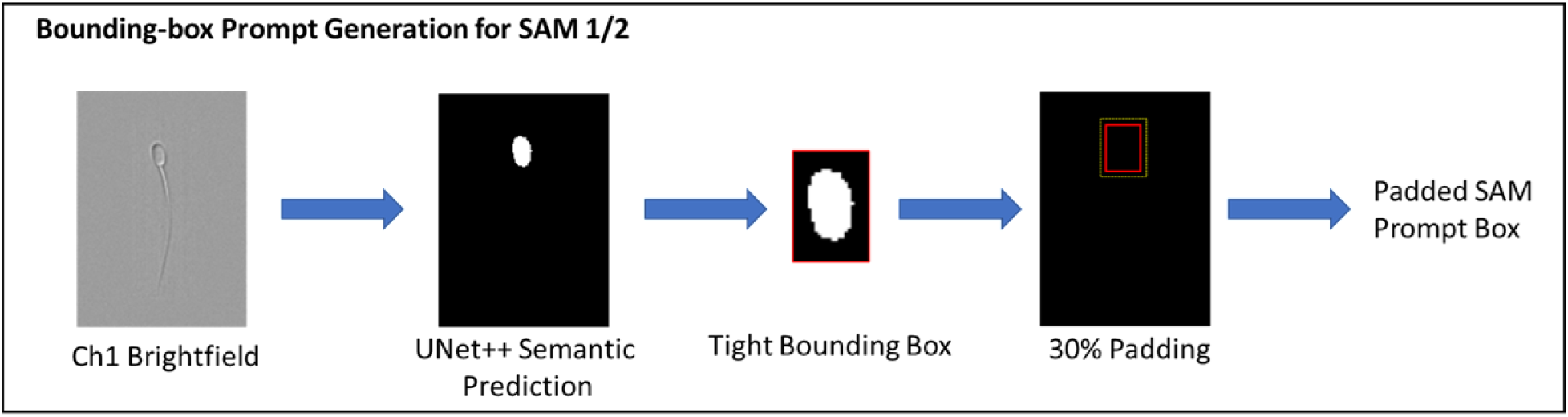
Bounding-box-prompted SAM refinement pipeline.

#### 2.5.2 ESRGAN Super-Resolution

To test whether native resolution imposed the accuracy ceiling, the Enhanced Super-Resolution Generative Adversarial Network was applied to upscale each brightfield image fourfold, from 165 × 120 to 660×480, and all five segmentation models were retrained on the upscaled data [21]. ESRGAN synthesizes high-frequency detail by optimizing a composite objective combining a pixel-fidelity term, a perceptual feature-space term, and a relativistic adversarial term, the last of which drives the generator to produce texture statistically consistent with natural high-resolution images [21].

Whether that synthesized detail helps is not obvious in advance. The label here is a macro-scale spatial support rather than a texture, and hallucinated content can introduce structural artifacts that decouple semantic boundaries from real biological features [22]. The setup is described here; Section 4 takes up the result.

### 2.6 Vision–Language-Model Head and Tail Delineation

#### 2.6.1 Gemma-4 Bounding-Box Annotation with SAM 2

Gemma-4 26B MoE was used to emit head and tail bounding boxes directly from the brightfield crop, which were then supplied to SAM 2 as box prompts [23, 24]. The model receives the crop alongside a system prompt that assigns it the role of a microscopy image-analysis assistant, defines sperm anatomy in terms of head, midpiece, and tail, supplies a constrained vocabulary of morphological defects, and requires a valid JSON object containing fractional bounding-box coordinates rather than prose [25]. The constrained output format is what makes the route programmatically usable at scale; the full protocol is given in Algorithm 1. Gemma-4 returns a box (x_1_, y_1_, x_2_, y_2_) for each region, and SAM 2 segments conditioned on it. Unlike the box derived from a UNet++ prediction in Section 2.5.1, this route localizes both the head *and* the tail from the brightfield image alone, and depends on neither the supervised network nor the fluorescence channel.

**Algorithm 1** Prompting protocol for automated head and tail annotation from brightfield imagery.

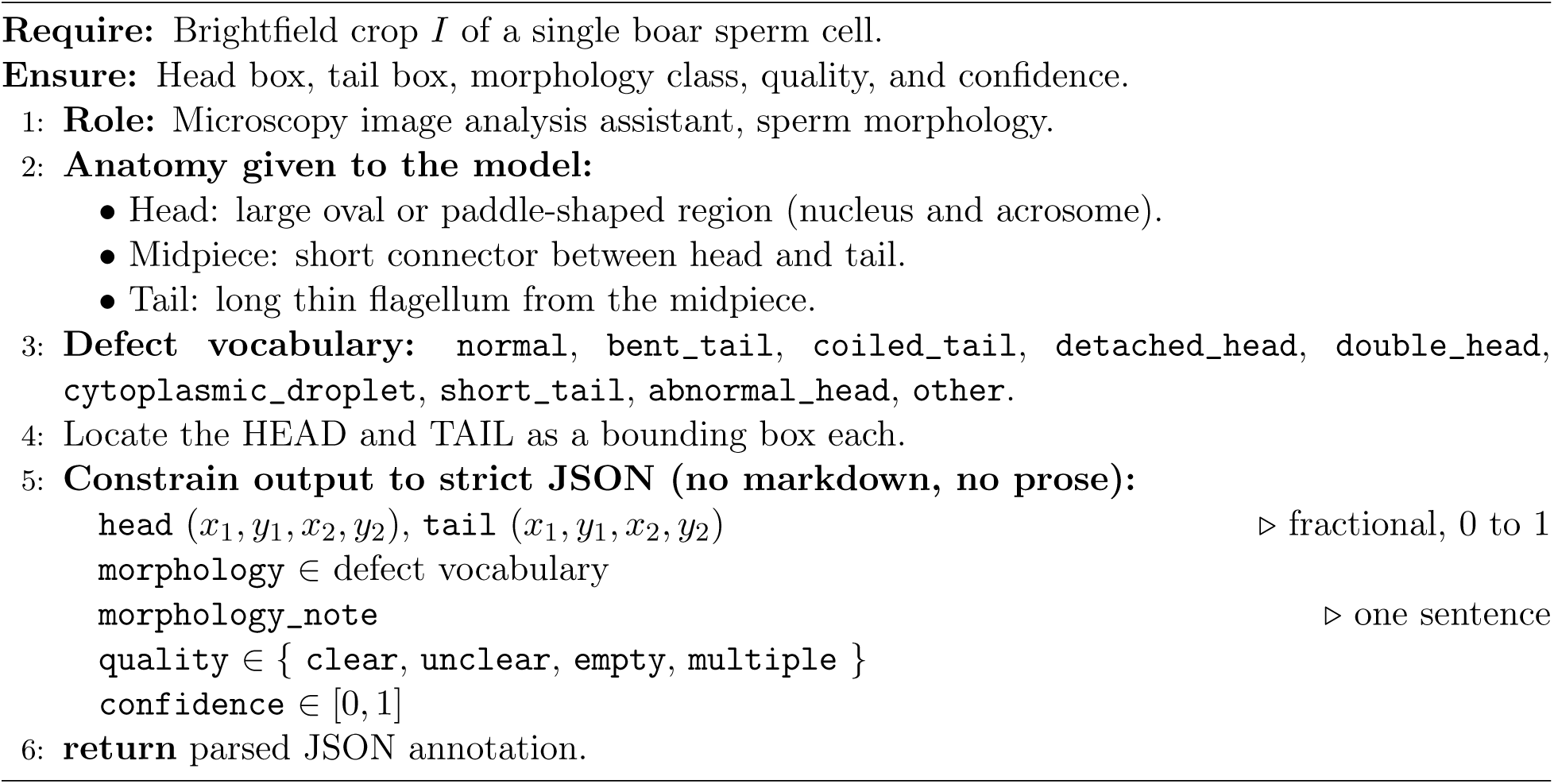

#### 2.6.2 Channel-Alignment Correction

The ImageStream instrument occasionally exhibits intrinsic channel-alignment errors, which surfaced when model predictions scored poorly against Ch7 despite tracking the visible cell closely: the two channels had drifted apart (Fig. 4). Such pairs must be identified before any pixel-level comparison between brightfield prediction and fluorescence reference is meaningful. Because Gemma-4 locates the head and tail from the brightfield image alone, its annotations are independent of the fluorescence channel and provide a reference against which the alignment of each Ch1/Ch7 pair can be assessed. The capability extends beyond the immediate task: vision–language annotations can serve as a diagnostic for instrument-level registration error, a point taken up in Section 4.

**Figure 4:**
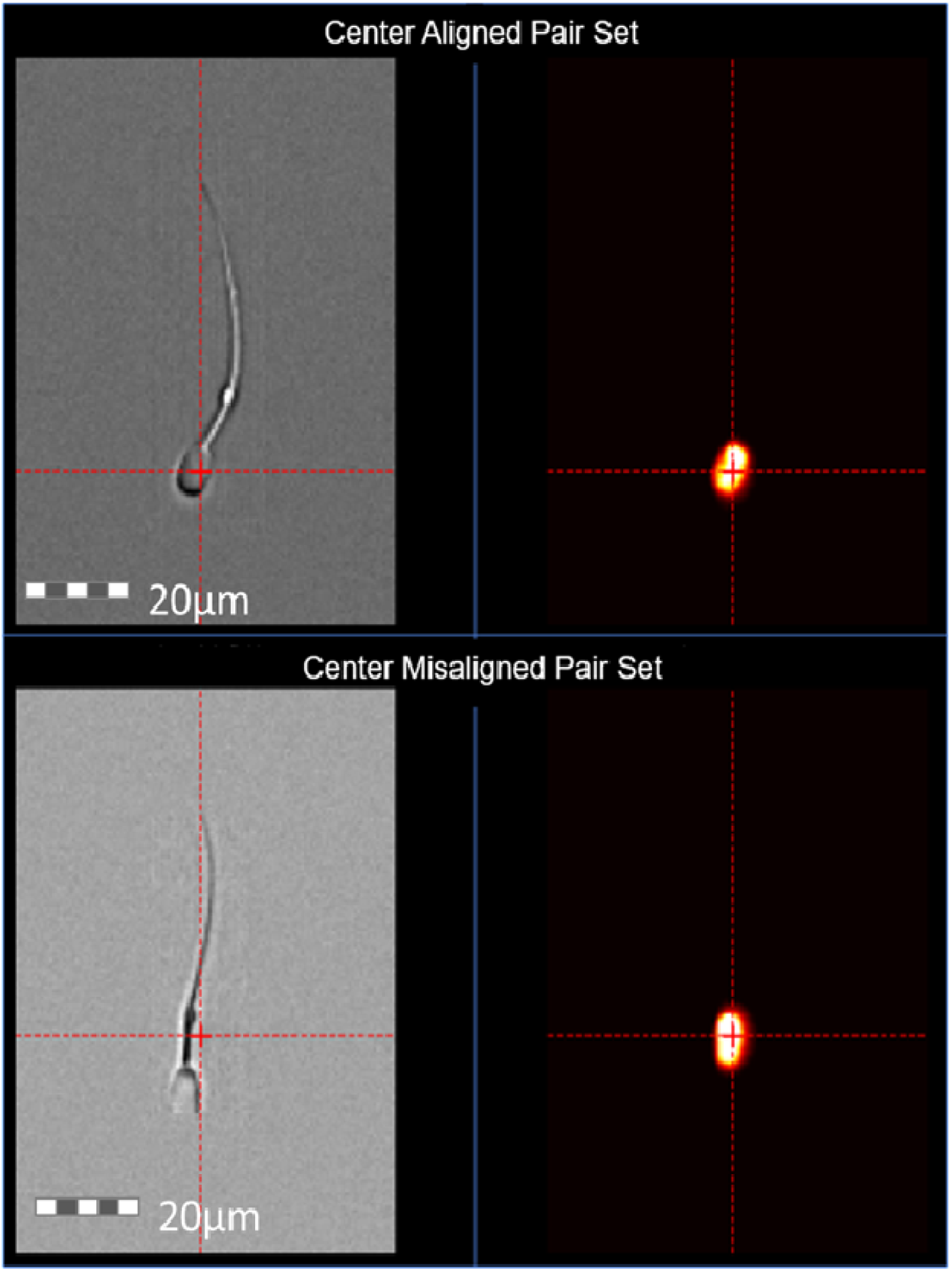
Example of correct channel alignment (top) versus an intrinsic alignment error (bottom).

#### 2.6.3 Construction of the Head-and-Tail Dataset

The annotation protocol was applied to the cleaned 11,420 image pairs. Cleaning proceeded by flagging every pair on which the supervised model scored below a Dice of 0.80, a threshold set just above the two-standard-deviation point of 0.792. All 712 flagged pairs were inspected for Ch1/Ch7 alignment, where misalignment means the fluorescence head had drifted up, down, left, or right relative to the brightfield head. The 312 pairs confirmed as misaligned were removed from the dataset.

The resulting data supports two parallel training branches, one on ESRGAN-upscaled imagery and one at native resolution, isolating the contribution of super-resolution within this regime. For direct comparison against the Hoechst 33342 reference, every generated label image was reduced to a binary membership map, since the quantity of interest is class membership (head, tail, or background) rather than pixel intensity.

#### 2.6.4 Three-Way Comparison

Three sources of segmentation are compared under a common protocol: a model trained on the Ch7 fluorescence label (the 11,420-pair dataset with the τ = 10 head mask), the same model refined by box-prompted SAM 2, and the Gemma-4 route applied zero-shot. All three are assessed with automated overlap metrics against the Ch7 reference, supplemented by human adjudication of whether the predicted mask is faithful at the pixel level. The comparison serves two purposes: it establishes how far a route that never sees the dye falls from one supervised by it, and it identifies the cells on which the two disagree, which Section 2.6.2 uses as a signal for channel misalignment. Results are reported in Section 3.

## 3 Results

This section reports the quantitative and qualitative outcomes of the five experimental stages. Interpretation is deferred to Section 4; what follows is a record of what was measured.

### 3.1 Classical Pipeline Ceilings

The erode–dilate–erode parameter sweep reached a hard ceiling. Across all schedule and threshold combinations tested, the Dice coefficient never exceeded 0.551, with that peak achieved under a 4–7–7 schedule at an adaptive threshold of T = 22. The four-stage head-hunting pipeline, which added geometric constraints, performed no better: a mean IoU of 0.31 and a mean Dice of 0.45 (Fig. 5).

**Figure 5:**
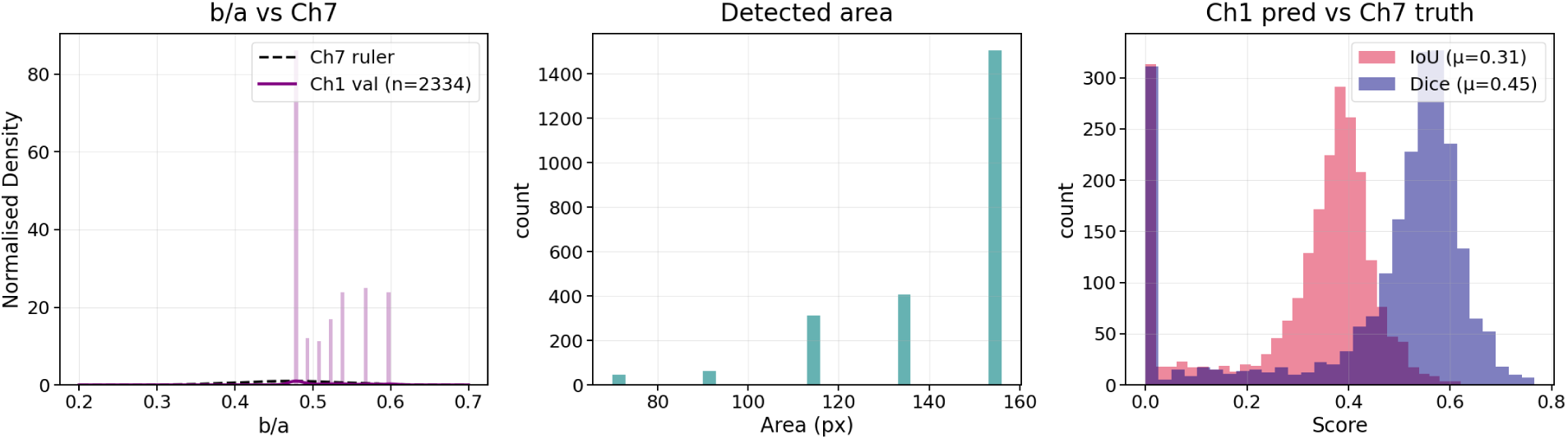
Four-stage classical pipeline performance. Left: distribution of detected b/a ratio against the Ch7 reference. Center: detected head area over the validation set. Right: per-cell Dice and IoU against the Ch7 ground truth.

The score distribution was strongly bimodal rather than uniformly mediocre. A substantial fraction of cells scored near zero, corresponding to complete localization failures, while the remainder clustered around 0.5. The near-zero population coincided with the smallest detected areas, indicating that the pipeline missed the head entirely on these events rather than misestimating its boundary.

The fluorescence-defined head aspect ratio spanned 0.31 to 0.60 across the dataset. Fixed-parameter operations localized the head acceptably on normal, well-oriented cells; on qualitative review, the near-zero failures corresponded to atypical morphology and low-contrast brightfield imagery.

### 3.2 U-Net-Based Models

Deep supervised networks resolved the morphological-variance limitations observed in the classical pipelines. Removing the 312 misaligned image pairs from the training dataset improved performance for the baseline configuration: UNet with a ResNet-50 encoder rose from a Dice of 0.9285 on the raw set to 0.9380 on the cleaned set. Because the misaligned pairs place the Ch7 head label off the true head, removing them raised recall (0.9348 → 0.9430) more than precision (0.9252 → 0.9354), consistent with the network recovering head pixels it had previously been penalized for predicting. All subsequent models were therefore trained on the cleaned dataset.

Table 2 reports the five architectures. The results are tightly clustered: every configuration lands within 0.6 percentage points of Dice (0.9343 to 0.9400), indicating that once the data is clean, this task is not architecture-limited. All five decoders track the head, and the differences between them are second-order refinements rather than categorical gaps. Two axes of variation nonetheless produce a consistent ordering.

**Table 2:**
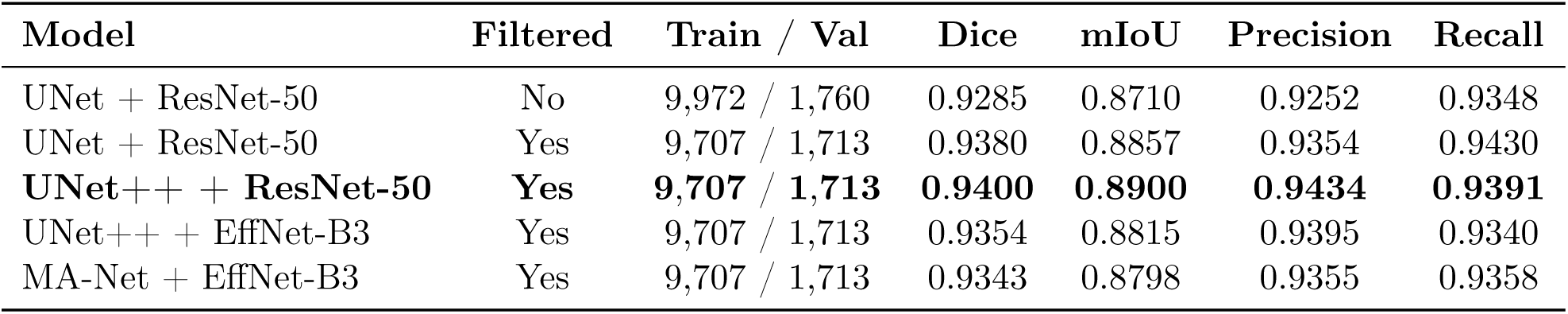
Segmentation performance across the five configurations. Filtered denotes removal of the 312 misaligned pairs.

| Model | Filtered | Train / Val | Dice | mIoU | Precision | Recall |
| --- | --- | --- | --- | --- | --- | --- |
| UNet + ResNet-50 | No | 9,972 / 1,760 | 0.9285 | 0.8710 | 0.9252 | 0.9348 |
| UNet + ResNet-50 | Yes | 9,707 / 1,713 | 0.9380 | 0.8857 | 0.9354 | 0.9430 |
| **UNet++ + ResNet-50** | **Yes** | **9,707 / 1,713** | **0.9400** | **0.8900** | **0.9434** | **0.9391** |
| UNet++ + EffNet-B3 | Yes | 9,707 / 1,713 | 0.9354 | 0.8815 | 0.9395 | 0.9340 |
| MA-Net + EffNet-B3 | Yes | 9,707 / 1,713 | 0.9343 | 0.8798 | 0.9355 | 0.9358 |

The first axis is the decoder family. Holding the ResNet-50 encoder fixed, replacing the plain UNet decoder with UNet++ raised Dice from 0.9380 to 0.9400 and mIoU from 0.8857 to 0.8900. UNet++ nests dense skip connections at multiple resolutions between encoder and decoder, so features are fused across several scales rather than passed through a single skip per stage. For an object like the sperm head, with almost all of its information concentrated on a short boundary, this multi-scale fusion helps resolve the exact edge, which is where the Dice score is won or lost. The gain is modest precisely because the head is a simple convex shape: the added skip pathways refine the boundary but cannot manufacture information the encoder did not already capture.

The second axis is the encoder backbone. Pairing UNet++ with the lighter EfficientNet-B3 rather than ResNet-50 lowered Dice from 0.9400 to 0.9354, and MA-Net with the same EfficientNet-B3 backbone landed at 0.9343. The parameter-efficient backbone did not surpass the heavier ResNet-50 on this task, counter to the usual expectation that EfficientNet matches or beats ResNet at far lower cost. The likely reason is domain and scale: EfficientNet-B3’s ImageNet-pretrained features and aggressive downsampling are tuned for large natural-image objects, whereas the head spans only 12–15 pixels at native resolution. ResNet-50’s simpler, higher-capacity feature stack appears to transfer more readily to this small, low-texture, single-channel target, where the discriminative signal is a faint intensity edge rather than the rich texture EfficientNet is optimized to exploit.

A finer point is visible in the precision–recall balance. The best model, UNet++ with ResNet-50, is the most precision-leaning of the cleaned configurations, meaning its predicted head pixels are the most reliable of the five even though its recall is not the highest: the cleaned UNet/ResNet-50 baseline posts a marginally higher recall (0.9430). In a downstream pipeline that uses head localization to seed tail recovery, precision-leaning behavior is the more useful failure mode, since a slightly conservative but trustworthy head boundary is preferable to an over-extended one that bleeds into the tail.

### 3.3 Foundation Models and Super-Resolution

#### 3.3.1 Zero-Shot and Prompted SAM Refinement

Applying the Segment Anything Model (SAM) as a refinement step over the UNet++ predictions alone lowered the measured head segmentation accuracy (Table 3). Prompted with the UNet++ bounding box expanded by a 30% margin, SAM1 (ViT-H) yielded a validation Dice of 0.707, acting conservatively (precision 0.895, recall 0.593). SAM2 (Hiera-Large) yielded a Dice of 0.743, acting more generously (precision 0.720, recall 0.790).

**Table 3:**
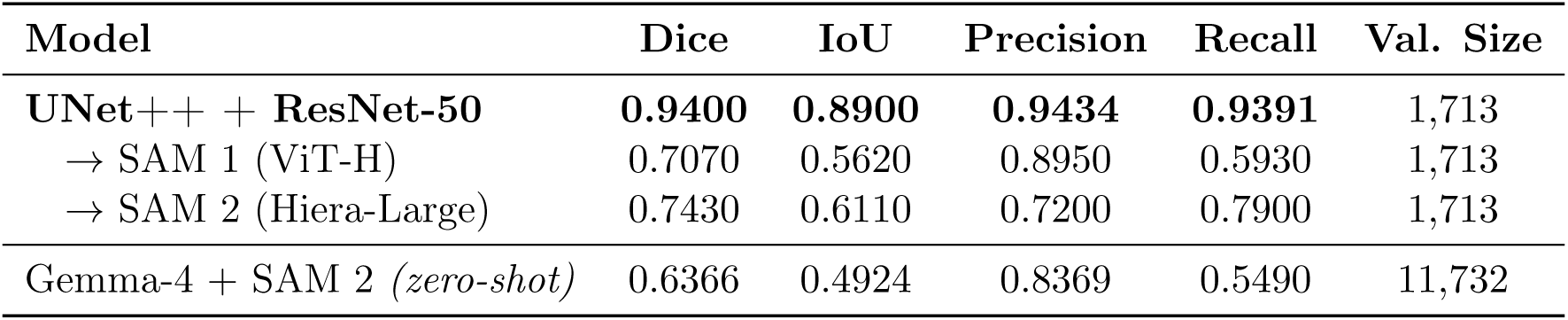
Zero-shot refinement and annotation results. Scores are measured against the Ch7 head reference, which does not label the tail. The zero-shot row is evaluated over the full uncleaned set and is therefore not directly comparable to the cleaned validation split used above it. The upper group reports the best supervised model and its box-prompted SAM refinement on the cleaned validation split; the lower group reports the zero-shot Gemma-4 + SAM 2 pipeline.

| Model | Dice | IoU | Precision | Recall | Val. Size |
| --- | --- | --- | --- | --- | --- |
| **UNet++ + ResNet-50** | **0.9400** | **0.8900** | **0.9434** | **0.9391** | 1,713 |
| → SAM 1 (ViT-H) | 0.7070 | 0.5620 | 0.8950 | 0.5930 | 1,713 |
| → SAM 2 (Hiera-Large) | 0.7430 | 0.6110 | 0.7200 | 0.7900 | 1,713 |
| Gemma-4 + SAM 2 (*zero-shot*) | 0.6366 | 0.4924 | 0.8369 | 0.5490 | 11,732 |

These scores warrant careful interpretation. The Dice and IoU are computed against the Ch7 fluorescence channel, which labels the chemically defined nuclear region rather than the full morphological cell. SAM1 and SAM2, operating purely on the brightfield Ch1 image, instead localize the visible cell body and edge. Part of the apparent accuracy drop therefore reflects a boundary-definition mismatch rather than a segmentation failure: both models often trace the cell competently but are penalized for not matching the smaller, stain-defined head target (Fig. 6).

**Figure 6:**
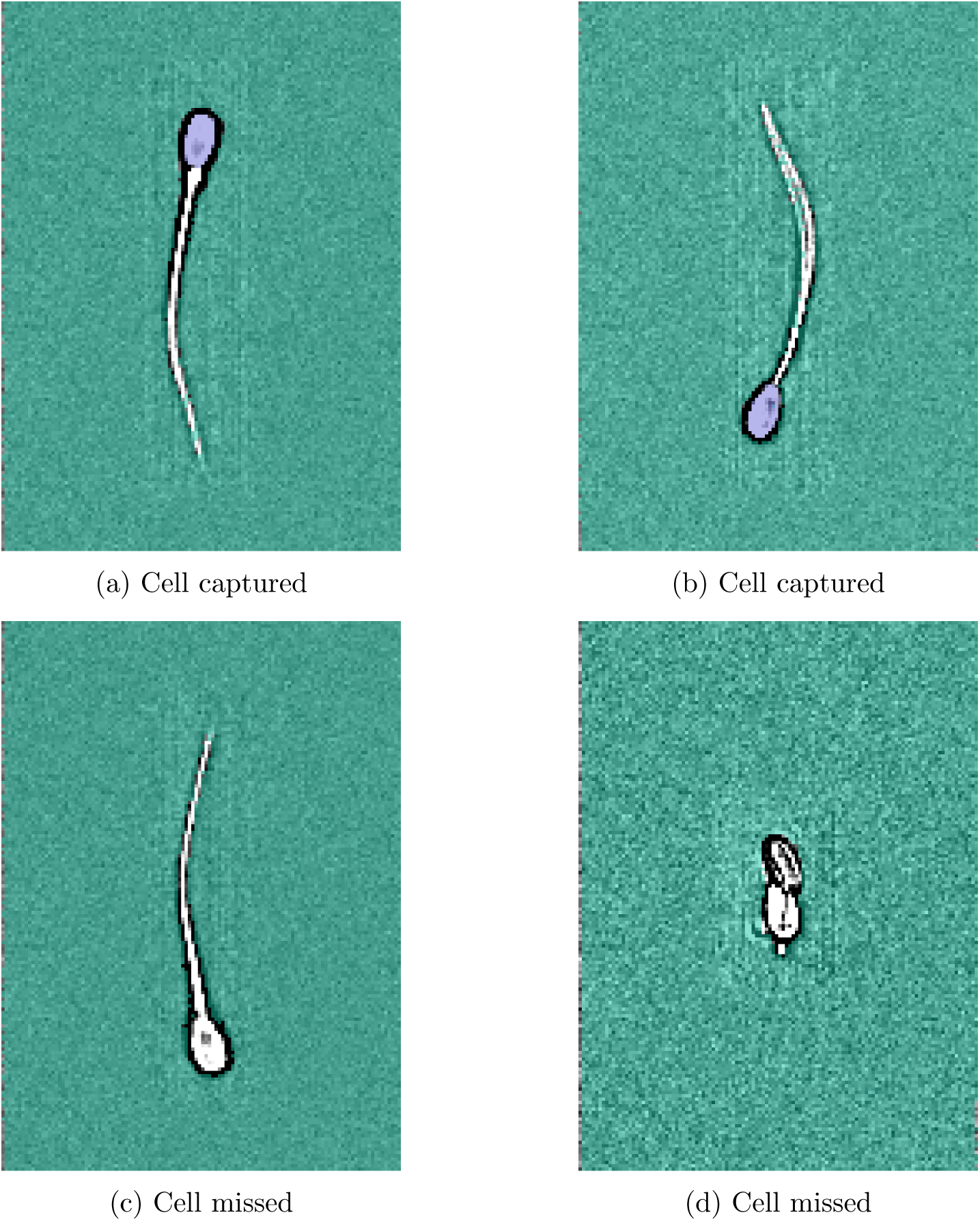
Representative Semantic Segmentation model + SAM2 outputs on Ch1 brightfield. Panels (a) and (b) show competent full-cell tracing, where SAM localizes the morphological cell head. Panels (c) and (d) show the failure mode, where the mask collapses onto background texture or a fragment of the cell and produces almost no overlap with the Ch7 head-only target.

The mismatch does not fully account for the low scores, however. Both models also miss the cell entirely on some events, latching onto background texture and producing near-zero overlap. Because these cases collapse the per-image score to zero, they depress the mean disproportionately and constitute a genuine reliability limitation independent of the metric mismatch.

#### 3.3.2 Gemma4-Prompted SAM2 Pipeline

Unlike the UNet++ bounding-box refinement, which merely re-segments a region the supervised network has already found, this pipeline is genuinely zero-shot: neither model is trained on the present data, and the head is located from scratch. Over the full validation set, the pipeline achieved a Dice of 0.6366 and an IoU of 0.4924 (Table 3), exceeding every classical handcrafted baseline while remaining below the supervised networks.

The error structure is more informative than the aggregate score suggests. High precision (0.8369) against lower recall (0.5490) indicates that when SAM 2 segments a region, that region is usually correct, but that it recovers only part of the head. Panel-level inspection attributes this to the prompting stage rather than the segmentation stage (Fig. 7). Where Gemma-4 places its box on the head, SAM 2 fills it accurately and overlap with the Ch7 reference is high (Dice 0.828). Where the box is displaced or undersized, SAM 2 segments faithfully within a region that contains little of the head, and the score collapses (Dice 0.039). Both channels are correctly registered in these examples, so the failure is distinct from the misalignment artifacts of Section 2.6.2.

**Figure 7:**
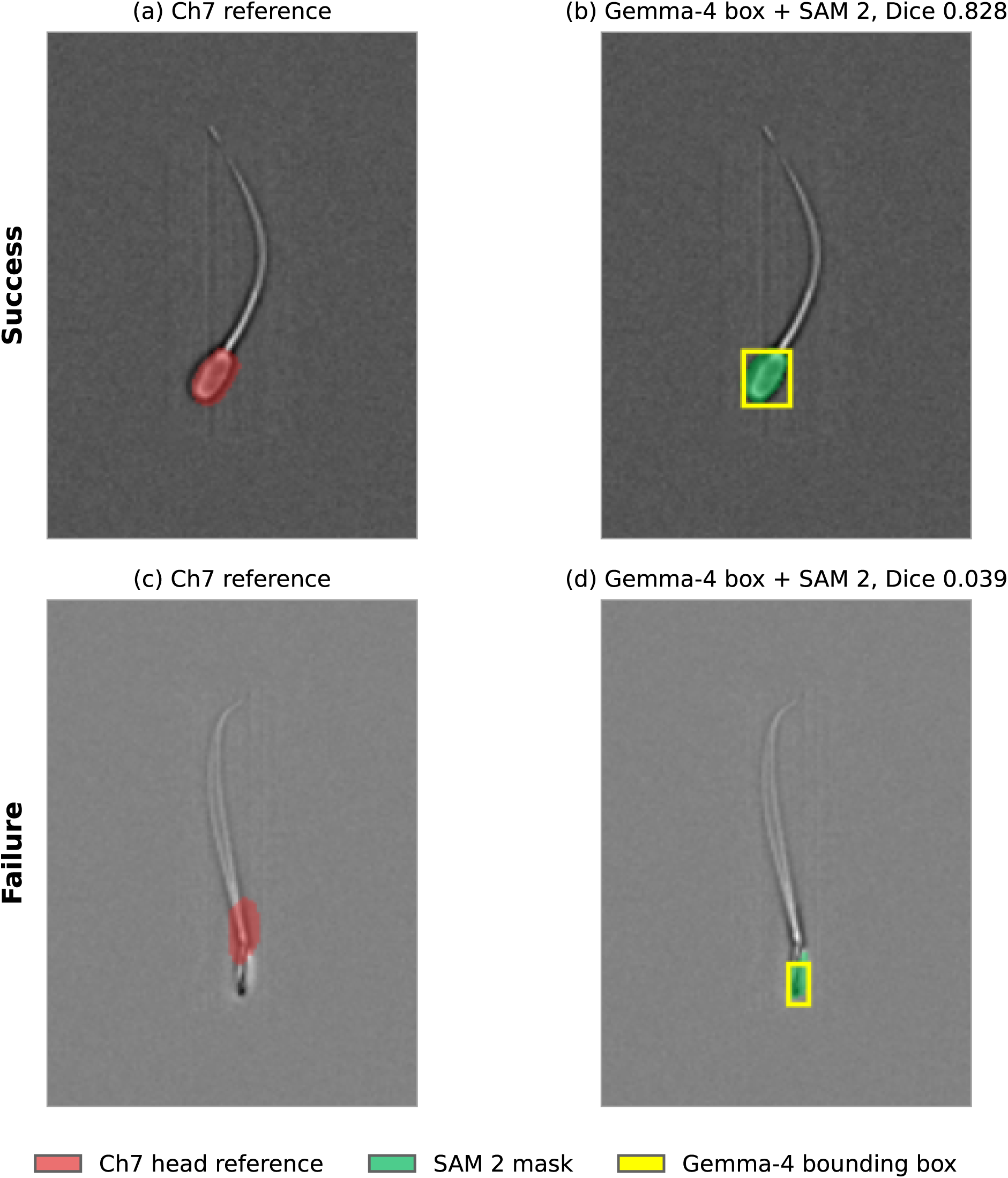
Zero-shot Gemma-4 + SAM 2 on two correctly registered pairs. Top: the box lands on the head and overlap is high. Bottom: the box is displaced and undersized, and overlap collapses. SAM 2 fills the box faithfully in both cases.

#### 3.3.3 ESRGAN Super-Resolution

Retraining the supervised configurations on the 4× ESRGAN-upscaled dataset produced a negligible absolute change in the Dice coefficient of +0.0025 (from 0.9400 to 0.9425). The four-fold artificial increase in pixel count did not translate into a meaningful gain in macro-scale spatial segmentation, confirming that native resolution is not the primary accuracy bottleneck for this dataset.

## 4 Discussion

The problem this work addresses is that every route to head and tail delineation at production scale has carried a disqualifying cost: manual tracing does not scale, fixed-parameter operators cannot absorb morphological variation, and supervised segmentation depends on a stain that is slow to apply, incapable of marking the flagellum, and silent about its own registration errors. What follows takes each of these in turn and examines how the two-route pipeline addresses it.

### 4.1 The Necessity of Learned Priors

Classical pipelines failed because they encode fixed priors. Rigid contrast gates and bounded aspect ratios cannot adapt to morphological variance: the head aspect ratio spans 0.31 to 0.60 across the dataset, and no single geometric parameterization covers that population without severe shape distortion. The ceiling of 0.551 Dice reported in Section 3.1 is the practical consequence.

Learned per-pixel models resolve this. A high-variance distribution favors data-driven adaptability over hand-specified rules [10, 26], which aligns with broader biological imaging domains where cellular plasticity has driven the same shift [27, 28]. The supervised network reaches 0.940 Dice from brightfield alone, and because the Ch7 label is consumed at training time only, no staining is required at inference.

### 4.2 Foundation Models: Domain Mismatch and Tail Labeling

Box-prompted SAM lowered the measured head Dice to 0.707 (SAM 1) and 0.743 (SAM 2). Two factors explain this. The first is a mismatch in target semantics: SAM is trained on natural images to segment salient boundaries such as edges and color transitions [13, 24], whereas the Hoechst 33342 reference marks fluorophore binding to the nuclear region. Fluorescence bloom offsets that binding from the visible brightfield boundary, so an accurate segmentation of the cell is scored as an error against a smaller target, and a one-to-three-pixel offset is a large relative error on a 150-pixel head. The second is localization: when the prompt drifts onto the tail or the background, SAM segments the wrong structure and overlap collapses. Much of the aggregate drop is therefore a boundary-definition artifact rather than a segmentation failure, consistent with evaluations showing that SAM struggles with low-contrast biomedical boundaries without fine-tuning [29, 30].

The same edge bias is what makes the route useful for the tail. Because Hoechst 33342 binds DNA confined to the condensed nucleus, the fluorescence protocol cannot mark the flagellum, so any tail supervision must come from a model that segments the whole cell. Subtracting the supervised head mask from a wide-prompted SAM 2 whole-cell mask would yield a computational tail label. We propose this as a route to tail pseudo-labels rather than a validated one: no hand-annotated tail set exists in this study, so its accuracy remains to be established.

### 4.3 The Signal-Processing Limits of Super-Resolution

The 4× ESRGAN upscale yielded a negligible Dice change of +0.0025. Super-resolution and macro-scale segmentation operate on disjoint spatial bands. ESRGAN minimizes perceptual objectives to synthesize high-frequency texture [21], whereas segmentation is a support-overlap task governed by low-frequency properties: position, area, and elongation. Upscaling adds no new low-frequency spatial support and therefore no usable information, consistent with the perception–distortion tradeoff [31]. Algorithms optimized for human perception do not by construction improve geometric reconstruction.

### 4.4 Vision–Language Annotation as an Independent Reference

Beyond the structures it cannot mark, the fluorescence protocol is also silent about its own failures: when the channel drifts out of registration, the result is a low score with no indication of why. Gemma-4 addresses this by annotating head and tail directly from brightfield [23]. Operating independently of the fluorescence optical path, its output serves as a channel-agnostic reference against which registration can be assessed, which is what made the 312 misregistered pairs identifiable at all. The recall-dominated recovery reported in Section 3.2 confirms that these were reference errors rather than model errors. An error mode previously invisible to the standard protocol therefore becomes a flagged, reviewable event, and the same mechanism would extend to any multi-channel acquisition where one channel is used to supervise another. This advances the shift toward in-silico labeling, in which algorithmic predictions supplement physical fluorophores where those fluorophores cannot reach [32, 33].

### 4.5 A Proposed Agreement-Based Routing Scheme

The foundation-model pathway scores below the supervised network against Ch7, but discarding it on that basis would waste the property that makes it useful: it fails on different cells. The results instead support combining the two, with the zero-shot pathway serving as an independent check (Fig. 9).

**Figure 8:**
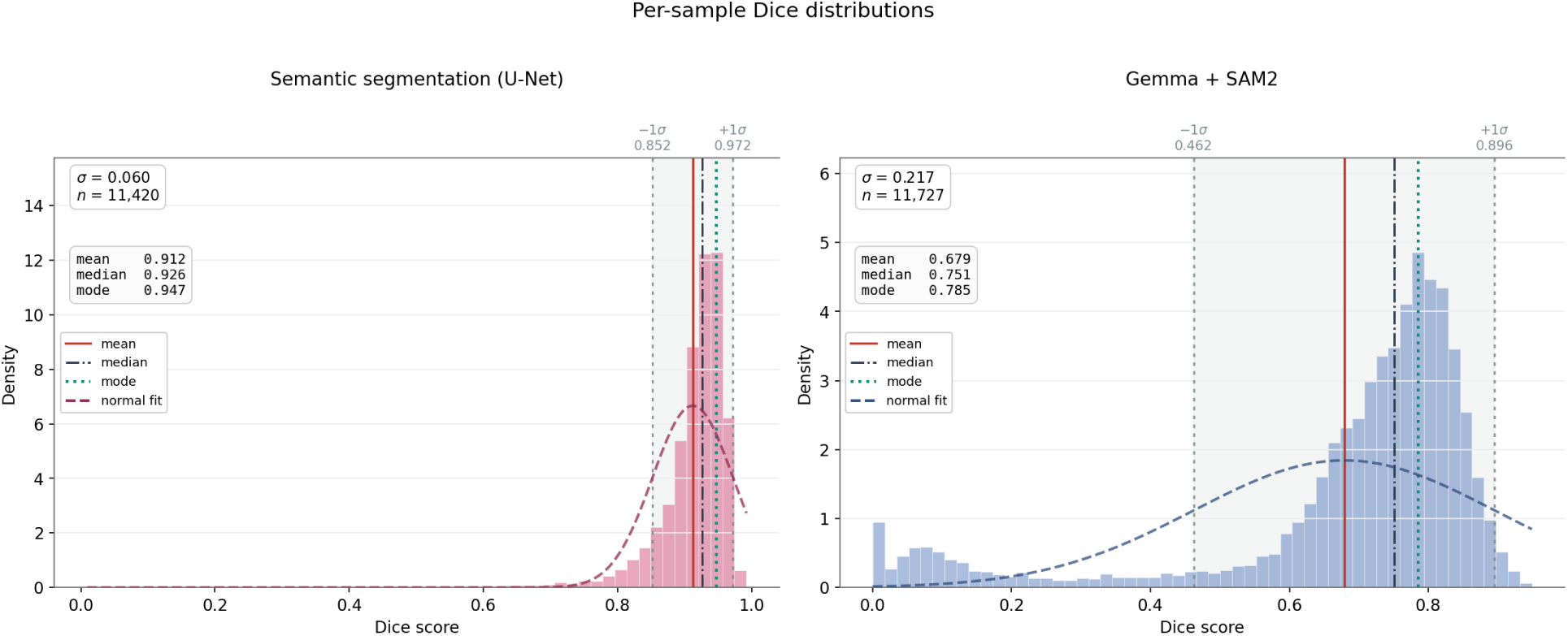
Per-sample Dice distributions for the two independent pathways. The supervised UNet network (left) is tightly concentrated near ceiling (mean 0.912, σ = 0.060), whereas the zero-shot Gemma-4 + SAM2 pathway (right) is broader and left-skewed (mean 0.679, σ = 0.217), with a distinct near-zero failure population. The two pathways fail on different cells, which is what makes their disagreement a useful anomaly signal.

**Figure 9:**
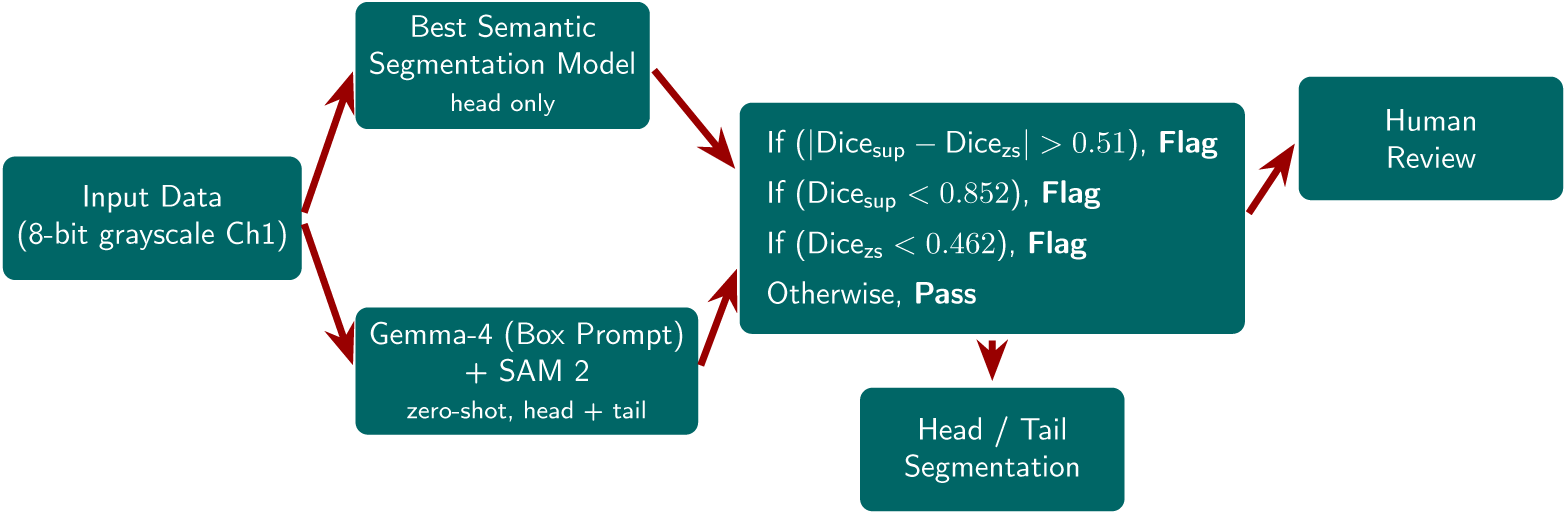
Proposed agreement-based routing. Both branches segment from Ch1 alone; cells on which they disagree, or on which either scores below its own −1σ point, are routed for human review rather than accepted.

The two pathways exhibit markedly different score distributions (Fig. 8). The supervised network is tightly concentrated near ceiling (mean Dice 0.912, σ = 0.060, computed per cell over all 11,420 cleaned pairs rather than on the held-out split of Table 2), while the zero-shot Gemma-4 + SAM 2 pathway is considerably broader (mean 0.679, σ = 0.217) and carries a distinct near-zero failure population. Because the two rest on different priors and fail on different cells, disagreement between them carries information about whether a given event is atypical.

This suggests a routing scheme, which we propose here rather than evaluate. Both pathways process the same Ch1 input independently. Rather than hand-tuning the gates, each threshold is taken from the one-standard-deviation band of the corresponding distribution, so that only events outside a pathway’s normal operating range are flagged. A cell would be routed to human review if any of the following holds:

- **High disagreement:** the absolute difference in Dice between the two pathways exceeds 0.51, the separation between the upper 1σ edge of the supervised distribution (0.972) and the lower 1σ edge of the zero-shot distribution (0.462). A gap wider than the two bands combined indicates the pathways genuinely disagree.
- **Low supervised confidence:** the supervised Dice falls below its own −1σ point of 0.852.
- **Low zero-shot confidence:** the Gemma-4 + SAM 2 Dice falls below its own −1σ point of 0.462.

The three gates catch different failures. Disagreement identifies cells where one pathway succeeds and the other does not, the signature of an out-of-distribution morphology or a prompt-placement error. The two low-confidence gates catch the case where both pathways score poorly against the same reference, which is what a misaligned pair produces, since a displaced label penalizes both routes at once. If none of the conditions is met, the cell passes and the automated head and tail masks are returned. Each gate corresponds to a departure beyond one standard deviation of expected behavior, so the scheme would preserve the throughput of the supervised network on standard data while routing atypical events to a human expert. Quantifying the flagging rate of each gate, and the precision of the resulting triage, is left to future work.

### 4.6 Limitations

Four limitations bound the claims made here. First, the reference is defined by where the fluorophore emits rather than where the cell boundary lies, so every Dice figure in this work is scored against an operational rather than an anatomical target; quantifying the exact offset requires a hand-annotated holdout set that this study does not provide. Second, no quantitative tail validation exists. The Gemma-4 route produces tail masks, and they are shown qualitatively, but establishing their accuracy requires pixel-level tail annotation that was not performed. Third, the exclusion of the 312 misaligned pairs depended on expert adjudication of 712 threshold-flagged cells, so the cleaning step is not reproducible from the threshold alone; the 400 cells that were flagged but not confirmed were retained. Fourth, the head label rests on a fixed intensity threshold (τ = 10), and any binary thresholding of a continuous fluorescence signal imposes a boundary the signal itself does not sharply define.

### 4.7 Future Work

The segmentation capability established here is a means to a larger end: a quantitative morphometric fingerprint of each spermatozoon. Reliable head and tail masks make interpretable descriptors available at single-cell resolution, including head area, head symmetry, and the length of the flagellum, which can then be aggregated into per-sire distributions. There is precedent for the biological value of such measurements: mitochondrial sheath length relates to fertility traits in boars used for artificial insemination, with conception rate and litter size associating in opposite directions [34]. The midpiece itself is the clearest gap, since the present acquisition protocol does not label it.

On the methodological side, the vision–language stage invites a systematic study of prompt engineering, and of arrangements in which specialized vision–language and segmentation models are composed rather than used in isolation. Training separate models tuned to individual defect classes, rather than one general-purpose network, is a further extension.

## 5 Conclusions

Head segmentation of boar spermatozoa can be performed from brightfield imaging-flow-cytometry data without fluorescent staining at inference. A UNet++ network with a ResNet-50 encoder, trained on labels derived from the Hoechst 33342 channel but shown only brightfield, reaches a Dice coefficient of 0.940, against a ceiling of 0.551 for the best handcrafted pipeline on the same data. Preserving native resolution by zero-padding rather than downscaling accounts for 2.5 points of that result; 4× generative upscaling contributes nothing measurable.

A second route, in which a vision–language model places bounding boxes that SAM 2 then segments, scores lower against the fluorescence reference but does two things the supervised route cannot. It delineates the flagellum, which no DNA-binding stain can mark, and because it depends on neither the network nor the dye, it gives an independent view of each cell. That independence makes the reference itself testable: 312 pairs in which the fluorescence channel had drifted out of registration were exposed by disagreement between the two routes, a failure the standard protocol has no means to detect.

Segmentation ground truth therefore need not be purely chemical. For a production andrology laboratory this removes the cost and delay of staining from routine morphological assessment, and it extends to species for which dye protocols are unavailable or impractical at scale.

## Ethics Statement

Semen used in this study was excess from commercial industry production and was not collected for the purposes of this study; it is therefore exempt from Institutional Animal Care and Use Committee (IACUC) oversight at Iowa State University.

## Acknowledgments

We would like to thank Iowa State start-up funding for Professor Anwesha Sarkar for this research.

## References

[1] Scott Pitnick, David J. Hosken, and Tim R. Birkhead. Sperm morphological diversity. In Tim R. Birkhead, David J. Hosken, and Scott Pitnick, editors, Sperm Biology: An Evolutionary Perspective, pages 69–149. Academic Press, London, 2009.

[2] Dorota Banaszewska and Katarzyna Andraszek. Assessment of the morphometry of heads of normal sperm and sperm with the Dag defect in the semen of Duroc boars. Journal of Veterinary Research, 65(2):239–244, 2021. doi: 10.2478/jvetres-2021-0019.

[3] Ashley Keller and Karl Kerns. Deep learning, artificial intelligence methods to predict boar sperm acrosome health. Animal Reproduction Science, 247:107–110, 2022.

[4] Ashley Keller, Molly K. Maus, Ella Keller, and Karl Kerns. Deep learning classification method for boar sperm morphology analysis. Andrology, 13(6):1615–1625, 2025.

[5] Jim Cummins. Sperm motility and energetics. In Tim R. Birkhead, David J. Hosken, and Scott Pitnick, editors, Sperm Biology: An Evolutionary Perspective, pages 185–206. Academic Press, London, 2009.

[6] ISU Single-Cell Phenomics Center. Single-cell phenomics center laboratory resources. https://faculty.sites.iastate.edu/kkerns/single-cell-phenomics-center. Last accessed on July 8, 2026.

[7] Nobuyuki Otsu. A threshold selection method from gray-level histograms. *IEEE Transactions on Systems*, Man, and Cybernetics, 9(1):62–66, 1979. doi: 10.1109/TSMC.1979.4310076.

[8] Ying Yu, Chunping Wang, Qiang Fu, Renke Kou, Fuyu Huang, Boxiong Yang, Tingting Yang, and Mingliang Gao. Techniques and challenges of image segmentation: A review. Electronics, 12(5):1199, 2023. doi: 10.3390/electronics12051199.

[9] Fariba Shaker, S. Amirhassan Monadjemi, and Ahmad Reza Naghsh-Nilchi. Automatic detection and segmentation of sperm head, acrosome and nucleus in microscopic images of human semen smears. Computer Methods and Programs in Biomedicine, 132:11–20, 2016. doi: 10.1016/j.cmpb.2016.04.026.

[10] Olaf Ronneberger, Philipp Fischer, and Thomas Brox. U-Net: Convolutional networks for biomedical image segmentation. In Medical Image Computing and Computer-Assisted Intervention (MICCAI), pages 234–241. Springer, 2015.

[11] Zongwei Zhou, Md Mahfuzur Rahman Siddiquee, Nima Tajbakhsh, and Jianming Liang. UNet++: A nested U-Net architecture for medical image segmentation. IEEE Transactions on Medical Imaging, 39(6):1856–1867, 2019. doi: 10.1109/TMI.2019.2959209.

[12] Tianshi Fan, Geng Wang, Yi Li, and Hongbin Wang. MA-Net: A multi-scale attention network for liver and tumor segmentation. IEEE Access, 8:184852–184865, 2020. doi: 10.1109/ACCESS.2020.3029411.

[13] Alexander Kirillov, Eric Mintun, Nikhila Ravi, Hanzi Mao, Chloe Rolland, Laura Gustafson, Tete Xiao, Spencer Whitehead, Alexander C. Berg, Wan-Yen Lo, and Piotr Dollár. Segment anything. In Proceedings of the IEEE/CVF International Conference on Computer Vision (ICCV), pages 4015–4026, 2023.

[14] Tyler Weide, Kayla Mills, Ian Shofner, Matthew W. Breitzman, and Karl Kerns. Metabolic shift in porcine spermatozoa during sperm capacitation-induced zinc flux. International Journal of Molecular Sciences, 25(14):7919, 2024. doi: 10.3390/ijms25147919.

[15] Ian J. Shofner, Kayla Mills, Tyler Weide, Matthew W. Breitzman, and Karl Kerns. Zinc regulation of lipidome remodeling during boar sperm capacitation. Journal of Animal Science, 104:skag009, 2026.

[16] Jean Serra. Image Analysis and Mathematical Morphology. Academic Press, London, 1982.

[17] Robert M Haralick, Stanley R Sternberg, and Xinhua Zhuang. Image analysis using mathematical morphology. IEEE Transactions on Pattern Analysis and Machine Intelligence, 9(4): 532–550, 1987.

[18] Rafael C. Gonzalez and Richard E. Woods. Digital Image Processing. Pearson, New York, NY, 4th edition, 2018.

[19] Alex Krizhevsky, Ilya Sutskever, and Geoffrey E Hinton. Imagenet classification with deep convolutional neural networks. In Advances in Neural Information Processing Systems (NeurIPS), volume 25, pages 1097–1105, 2012.

[20] Oystein Haugen, Nooor Al-Sharhan, Michael A. Riegler, Pal Halvorsen, and Hugo L. Hammer. Ensembling noisy segmentation masks of blurred sperm images. Scientific Reports, 13(1):16602, 2023. doi: 10.1038/s41598-023-43501-y.

[21] Xintao Wang, Ke Yu, Shangxiang Wu, Jinjin Gu, Yihao Liu, Chao Dong, Yu Qiao, and Chen Change Loy. ESRGAN: Enhanced super-resolution generative adversarial networks. In Proceedings of the European Conference on Computer Vision (ECCV) Workshops, pages 0–0, 2018.

[22] Berta Marghalani, Muhammad Khan, and Shadi Mostafa. Generative adversarial networks for medical image super-resolution: A review of applications, opportunities, and challenges. Biomedical Signal Processing and Control, 89:105740, 2024. doi: 10.1016/j.bspc.2023.105740.

[23] Gemma Team, Thomas Mesnard, Cassidy Hardin, Robert Dadashi, Surya Bhupatiraju, Shayan Shakeri, Aakanksha Andreassen, et al. Gemma: Open models based on gemini research and technology. *arXiv preprint arXiv:2403.08295*, 2024.

[24] Nikhila Ravi, Valentin Gabeur, Yuan-Ting Hu, Ronghang Hu, Chaitanya Ryali, Tengyu Banna, et al. SAM 2: Segment anything in images and videos. *arXiv preprint arXiv:2408.00714*, 2024.

[25] Haotian Liu, Chunyuan Li, Qingyang Wu, and Yong Jae Lee. Visual instruction tuning. Advances in Neural Information Processing Systems, 36, 2024.

[26] Geert Litjens, Thijs Kooi, Babak Ehteshami Bejnordi, Arnaud Alban Setio, Francesco Ciompi, Mohsen Ghafoorian, Jeroen AWM van der Laak, Bram van Ginneken, and Clara I Sánchez. A survey on deep learning in medical image analysis. Medical Image Analysis, 42:60–88, 2017. doi: 10.1016/j.media.2017.07.005.

[27] Thorsten Falk, Dominic Mai, Robert Bensch, Özgün Çiçek, Ahmed Abdulkadir, Yassine Marrakchi, Anton Böhm, Jan Deubner, Zoe Jäckel, Katharina Seiwald, et al. U-net: deep learning for cell counting, detection, and morphometry. Nature Methods, 16(1):67–70, 2019.

[28] Carsen Stringer, Tim Wang, Michalis Michaelos, and Marius Pachitariu. Cellpose: a generalist algorithm for cellular segmentation. Nature Methods, 18(1):100–106, 2021.

[29] Maciej A Mazurowski, Haoyu Dong, Hanxue Gu, Jichen Yang, Nicholas Kurosu, et al. Segment anything model for medical image analysis: an experimental study. Medical Image Analysis, 89:102918, 2023.

[30] Wei Ji, Junde Li, Qi Bi, Wenbo Li, and Li Cheng. SAM-Med2D: Segment anything model for 2d medical imaging. *arXiv preprint arXiv:2308.16184*, 2023.

[31] Yochai Blau and Tomer Michaeli. The perception-distortion tradeoff. In Proceedings of the IEEE Conference on Computer Vision and Pattern Recognition, pages 6228–6237, 2018.

[32] Eric M Christiansen, Samuel J Yang, D Michael Ando, Ashkan Javaherian, Gaia Skandarajah, Scott Kelley, Hunter A Rubin, William A Lee, Michael D Carroll, Christophe Brenner, et al. In silico labeling: predicting fluorescent labels in unlabeled images. Cell, 173(3):792–803, 2018.

[33] Tao Tu, Shekoofeh Azizi, Danny Driess, Mike Schaekermann, Rohan Amin, Pi-Chuan Chang, Andrew Carroll, Chuck Lau, Ryutaro Tanno, Ira Ktena, et al. Towards generalist biomedical AI. NEJM AI, 1(3):AIoa2300138, 2024.

[34] Karl Kerns, Jennifer Jankovitz, Julie Robinson, Amanda Minton, Chris Kuster, and Peter Sutovsky. Relationship between the length of sperm tail mitochondrial sheath and fertility traits in boars used for artificial insemination. Antioxidants, 9(11):1033, 2020. doi: 10.3390/antiox9111033.

